# Reverse immunodynamics: a new method for identifying targets of protective immunity

**DOI:** 10.1101/159483

**Authors:** Katrina J Spensley, Paul S Wikramaratna, Bridget S Penman, Andrew Walker, Adrian L Smith, Oliver G Pybus, Létitia Jean, Sunetra Gupta, José Lourenço

**Affiliations:** Imperial College London, London W2 1PG; Institute of Evolutionary Biology, University of Edinburgh, Edinburgh EH9 3JT; Department of Zoology, University of Oxford, Oxford OX1 3PS; The Sir William Dunn School of Pathology, University of Oxford, Oxford OX1 3RE

## Abstract

Despite a dramatic increase in our ability to catalogue variation among pathogen genomes, we have made far fewer advances in using this information to identify targets of protective immunity. We propose a novel methodology that combines predictions from epidemiological models with phylogenetic and structural analyses to identify such targets. Epidemiological models predict that strong immune selection can cause antigenic variants to exist in non-overlapping combinations. A corollary of this theory is that targets of immunity may be identified by searching for non-overlapping associations among antigenic variants. We applied this concept to the AMA-1 protein of the malaria parasite *Plasmodium falciparum* and found strong signatures of immune selection among certain regions of low variability which could render them ideal vaccine candidates.

## Introduction

Apical membrane antigen 1 (AMA-1) is a trans-membrane protein common to all *Plasmodium* species with a putative role in invasion at erythrocytic and pre-erythrocytic stages of the parasite life-cycle. Antibodies to AMA-1 appear to inhibit erythrocyte invasion and are thus thought to play a role in clinical protection against both *P. falciparum* [1-7] and *P. vivax* [8]. Cellular responses to AMA-1 may provide immunity against both erythrocytic [9] and pre-erythrocytic forms of *P. falciparum* [10]. The potential of AMA-1 as a component of a multi-stage malaria vaccine is compromised by the polymorphism of many of the targets of natural immunity [11,12] as well as by difficulties in inducing lasting immune responses through vaccination [13-16]. Identifying variable regions of AMA-1 that are under strong immune selection would afford the possibility of focusing vaccine-induced immune responses on these regions and ensure full coverage of variability at these critical epitopes.

The extracellular region of AMA-1 can be structurally resolved into three domains: DI, DII and DIII. The highest level of polymorphism is seen among residues in DI and DII surrounding a conserved hydrophobic trough [17,18] which plays a vital role in the invasion process [19-21]. Antibodies against this region have been shown to block invasion in a strain-specific manner; however, it is also clear that these antibodies constitute only a small proportion of the total inhibitory humoral response [12,22,23]. *Pf*AMA1 also contains a number of less variable residues across the three domains, and our aim was to identify which of these were also located within epitopes eliciting protective immune responses.

We attempted to identify these additional epitopes under strong immune selection by using a novel approach based on the predictions of a multilocus mathematical model of pathogen evolution [24-26]. The central premise is that regions or epitopes under strong immune selection will self-organise into non-overlapping combinations of variants [24-26] and supported by empirical observations within a variety of host-pathogen systems [27-32]. We inverted this result to search for signatures of immune selection among polymorphic loci by identifying those pairs which exhibit a high degree of ‘non-overlap’ but excluding those that may have arisen as a consequence of shared ancestry or due to structural interactions. Using this “reverse immunodynamic” method, we found strong signatures of immune selection among a small number of dimorphic residues in Domain III of *Pf*AMA-1, suggesting that epitopes containing these residues are targets of protective immunity.

## Results

### A limited number of site-pairs exhibit high non-overlap

We implemented the reverse immunodynamic method by performing pairwise comparisons of all dimorphic sites among 1198 unique *Pf*AMA-1 sequences obtained from GenBank, to determine whether any of these pairs exhibited non-overlapping associations between amino acid residues. We used an Information Theory statistic known as Mutual Information (MI), scaled to its maximum value, to measure the degree of non-overlap in amino acid combinations between pairs of sites. Most pairs of sites showed low non-overlap, with only 10 pairs exhibiting a scaled MI in excess of 0.3 (Tables 1 & S1). Of these, pairs 404/405, 448/451 and 283/285 were eliminated on grounds of structural proximity, leaving 7 pairs as shown by the bars in Figure 1A.

**Table 1:** Frequencies of amino acid associations between pairs of sites with high nonoverlap (scaled MI>0.3)

| Association | Amino Acid Combinations and Frequencies |  |  |  | Measure of non-overlap (scaled MI) |
| --- | --- | --- | --- | --- | --- |
| **496 vs 503** | **IN** | **IR** | **MN** | **MR** | 0.89 |
|  | 602 | 3 | 14 | 561 |  |
| **503 vs 512** | **NK** | **NR** | **RK** | **RR** | 0.61 |
|  | 565 | 52 | 40 | 524 |  |
| **451 vs 485** | **KI** | **KK** | **MI** | **MK** | 0.57 |
|  | 604 | 103 | 15 | 476 |  |
| **496 vs 512** | **IK** | **IR** | **MK** | **MR** | 0.53 |
|  | 552 | 68 | 54 | 523 |  |
| **283 vs 285** | **LE** | **LQ** | **SE** | **SQ** | 0.52 |
|  | 242 | 71 | 1 | 883 |  |
| **439 vs 451** | **HK** | **HM** | **NK** | **NM** | 0.5 |
|  | 660 | 72 | 46 | 390 |  |
| **206 vs 225** | **EI** | **EN** | **KI** | **KN** | 0.5 |
|  | 96 | 806 | 290 | 6 |  |
| **439 vs 485** | **HI** | **HK** | **NI** | **NK** | 0.41 |
|  | 588 | 144 | 30 | 406 |  |
| **448 vs 451** | **DK** | **DM** | **NK** | **NM** | 0.39 |
|  | 705 | 196 | 2 | 295 |  |
| **404 vs 405** | **RE** | **RK** | **TE** | **TK** | 0.34 |
|  | 108 | 635 | 367 | 84 |  |

**Figure 1:**
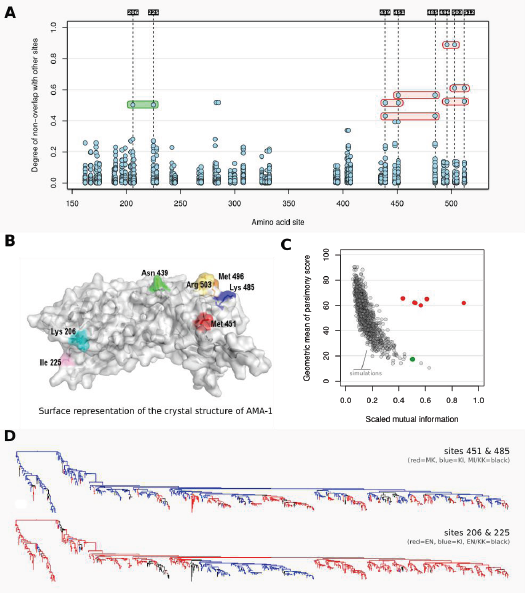
(A) Degree of non-overlap in associations between amino acid residues at pairs of dimorphic sites. Each site on the x-axis is compared with all other dimorphic sites: those with reciprocally high scores are shown by bars spanning both sites. Red bars show significant associations. Green bars represent associations that do not achieve significance when phylogenetic relationships are taken into account. (B) Surface representation of the crystal structure of AMA-1 with specific amino acids highlighted: Lysine (Lys) 206 = Cyan; Isoleucine (Ile) 225 = Magenta; Asparagine (Asn) 439 = Green; Methionine (Met) 451 = Red, Lys 485 = Blue, Met 496 = Orange, Arginine (Arg) 503 = Yellow. (C) Pairs of sites shown by red and green bars in Fig1A are shown in relation to a null distribution of site-pairs from 1000 simulations that scored most highly on the product of scaled Mutual Information (measuring non-overlap) and Geometric Mean of Parsimony Scores (indicating the minimum number of mutations required to explain the observed ancestral relationships). (D) Phylogenetic trees of AMA-1 with branches coloured on the basis of observed combination of amino acid variants found at tips for sites 451 and 485 (red = MK, blue = KI, grey = MI/KK) and sites 206 and 225 (red = EN, blue = KI, grey = EI/KN) using “Annotate Nodes from Tips” in FigTree. Black branches indicate nodes whose annotation cannot be resolved by the algorithm within FigTree.

### High non-overlap is unlikely to have arisen by neutral evolutionary processes

To assess whether high non-overlap between sites could have arisen by neutral evolutionary processes, we estimated a phylogenetic tree from the sequences and calculated the parsimony score (PS) for each site, which indicates the minimum number of mutations (in this case, back and forth between the two possible amino acids at each site) required to explain the observed ancestral relationships. A new statistic S, the product of MI and the geometric mean of PS for each site, was found to preserve the same order as the scaled MI scores of the 7 sites under consideration, except in the case of the 206/225 (Table S2). To obtain an appropriate null distribution of S for hypothesis testing, we simulated 1000 sets of sequences which were (a) consistent with the empirical phylogenetic tree and (b) had the same PS score at each dimorphic site as the empirical data. The null distribution of the top-scoring measures of S (Figure 1C) shows that neutral evolution can generate sites with high MI or high PS values, but not both. In contrast, the pairs of sites indicated in red in Figure 1A lie well outside the null distribution of high-scoring site-pairs (Figure 1C). Only site-pair 206/225 (shown in green) did not meet this criterion. Reasons for this pattern become evident upon inspection of the distribution of the amino acid variation on the phylogenetic tree (Figure 1D). The dominant non-overlapping pairs of amino acids at 206/225 are highly segregated on the phylogeny and not improbable under the null hypothesis, while those of 451/485, for example, exhibit a more interdigitated pattern, indicating multiple independent and convergent paired amino acid changes that are exceptionally unlikely in the absence of immune selection.

### Analysis of tertiary structure identifies those sites that are most likely to be under immune selection

The group of sites which scored most highly on the conservative test (described above) comprised 496, 503 and 512. However, mapping these onto the tertiary structure of AMA-1 (Figure 1B) shows that they are in very close proximity of each other and therefore we were unable to exclude biochemical interactions as the primary cause of this striking association. It is still possible that variation at these sites is held in strong non-overlap through immune selection. Antibodies specific to epitopes containing amino acids I/N/K may cross-react with all other epitopic variants except those containing M/R/R, thereby promoting the co-circulation of these non-overlapping combinations of residues at these sites; however, because of their structural adjacency, it is impossible to privilege this explanation over intrinsic steric advantages to these combinations.

High non-overlap was also observed among sites 439/451, 439/485 and 451/485 which were sufficiently far enough apart in the tertiary structure to represent potential examples of immune selection forcing non-structurally related residues to co-segregate. We cannot, of course, completely eliminate long-range physical interactions as a cause of these associations, and whether it is indeed immune selection that is responsible for these patterns can only be established by functional studies. It is clear that there is no fundamental structural impediment to the existence of intermediate forms since they do occur at low frequencies, but whether they are disadvantaged due to lower intrinsic fitness or immune-mediated selection remains to be resolved by further experimental work.

### Non-overlapping associations between sites under strong immune selection are observed at the level of local populations

We next investigated the associations between positions 439, 451 and 485 on a smaller geographical scale to confirm that these patterns did not arise due to certain combinations of alleles dominating by chance in particular areas. In all countries studied, both of the major sequence variant associations were present, with the secondary variants present at a lower frequency (Table 2).

**Table 2:** Country-specific frequency of amino acid associations between positions 439, 451 and 485 within unique sequences obtained from The Gambia, Kenya and Mali

| Association | Country | Amino Acid Combinations and Frequencies |  |  |  | Scaled MI |
| --- | --- | --- | --- | --- | --- | --- |
| **439 vs 451** |  | **HK** | **HM** | **NK** | **NM** |  |
|  | **Gambia** | 46 | 3 | 0 | 31 | 0.701 |
|  | **Kenya** | 46 | 3 | 9 | 27 | 0.402 |
|  | **Mali** | 395 | 25 | 11 | 211 | 0.637 |
| **439 vs 485** |  | **HI** | **HK** | **NI** | **NK** |  |
|  | **Gambia** | 39 | 7 | 1 | 30 | 0.549 |
|  | **Kenya** | 45 | 4 | 1 | 35 | 0.682 |
|  | **Mali** | 359 | 61 | 17 | 205 | 0.453 |
| **451 vs 485** |  | **KI** | **KK** | **MI** | **MK** |  |
|  | **Gambia** | 39 | 4 | 1 | 35 | 0.673 |
|  | **Kenya** | 46 | 9 | 0 | 31 | 0.515 |
|  | **Mali** | 366 | 40 | 10 | 244 | 0.608 |

A dataset of 660 unique sequences was compiled for Mali and showed high non-overlap for each pair of associations. A smaller dataset comprising 79 sequences from the Gambia also yielded MI values in excess of those obtained from the global data set. Slightly weaker associations were found in 86 unique sequences from Kenya; however these improved significantly when the data were analysed as a collection of isolates rather than sequences: i.e. without excluding identical sequences present in different individuals (Table 3).

**Table 3:** Frequency of amino acid associations between positions 439, 451 and 485 in a population study (duplicate sequences not excluded) in Kilifi, Kenya

| **Association** | **Amino Acid Combinations and Frequencies** |  |  |  | **Scaled MI** |
| --- | --- | --- | --- | --- | --- |
| **439 vs 451** | **HK** | **HM** | **NK** | **NM** | 0.564 |
|  | 61 | 3 | 9 | 55 |  |
| **439 vs 485** | **HI** | **HK** | **NI** | **NK** | 0.772 |
|  | 60 | 4 | 1 | 63 |  |
| **451 vs 485** | **KI** | **KK** | **MI** | **MK** | 0.644 |
|  | 61 | 9 | 0 | 59 |  |

As the model predictions pertain to population level frequencies of allele combinations, rather than their presence in a collection of unique sequences, we believe that Table 3 presents a more valid test (than when only unique sequences are included) of the model’s capabilities to pick out regions under strong immune selection; it is reassuring that these patterns are also evident under the more stringent conditions where only unique sequences are included.

## Discussion

Our results suggest that Domain III (DIII) of the AMA-1 protein is under strong immune selection, and may play an important role in protective immunity alongside the more polymorphic C1L region of Domain I. At a functional level, Domain III has been shown to bind to the erythrocyte membrane protein Kx [33]. Immune responses that interrupt this process may, therefore, have a significant inhibitory effect on parasite growth but, so far, the evidence that Domain III elicits protective antibody responses is limited. Nair *et al.* [34] have shown that antibodies from a single blood donor affinity purified on DIII are inhibitory. In a study by Polley *et al.* [2], it was found that antibody responses against recombinant full-length ectodomains were more strongly associated with protection than responses to those which did not express DIII, even though naturally acquired antibodies to recombinant DIII alone were rare. A virosomal formulation of loop I of DIII, comprising residues 446-490, has been shown to elicit growth-inhibitory antibodies [35] and volunteers within a vaccine trial containing this AMA-1 peptide exhibited both reduced blood stage growth rates and a higher rate of morphological parasite abnormalities [36]. A recent analysis of allele-specific efficacy of a vaccine against AMA-1 [11] showed that the relative risk ratio (RRR) of re-infection with strains containing vaccine-type amino acids among vaccinees vs controls was high (although not significant) at residues 439, 451 and 485. Residues associated with the polymorphic sites surrounding the hydrophobic trough displayed significant RRR, as might be expected; sites 308 and 332 also had high scores but no evidence of sustained immune selection at these sites was available in our analysis. In the same study, the likelihood of a shift from a vaccine-type to non-vaccine-type residue among sequences obtained following vaccination was found to be significantly increased at position 485 and non-significantly increased at position 451. Significant changes in residue composition were also found at highly polymorphic site 197 but not at any of the other polymorpic sites in the C1L region. Significant change was also observed at dimorphic site 435, but we found no non-overlapping associations in any of the pairwise comparisons between 435 and other sites. Several population genetic studies [37-41] based on site frequency spectrum analysis techniques (such as Tajima’s D) strongly indicate that DIII is under balancing selection; our “reverse immunodynamic” method complements and extends these analyses by showing that we can use a measure of non-overlap between site-pairs in addition to standard population genetic measures to finely map regions under immune selection within DIII.

An interesting feature of our analyses is that while particular groups of sites exhibit strong non-overlap, the associations between these groups appear to be random. For example, the triad 439-451-485 is in complete linkage equilibrium with triad 496-503-512. Furthermore, there are no non-overlapping associations present between the highly polymorphic residues in the C1L region of Domain I and the 439-451-485 triad (although the former exhibit strong non-overlap: Table S1). If the analysis we present here is correct, then it would be sufficient to present the dominant “non-overlapping” variants of 439-451-485 (i.e. HKI and NMK) in a manner that induced strong immune responses to the epitopes containing these residues, rather than to try and cover the entire haplotypic diversity of AMA-1 [42]. Immune responses to the epitopes containing residues 439, 451 and 485 appear to develop more slowly under natural conditions than antibodies to the highly polymorphic regions shielding the hydrophobic groove [2], but should be easier to induce by vaccination than immune responses to completely conserved epitopes [43].

Our analysis of AMA-1 further highlights the role of less polymorphic epitopes in population-level immune evasion. At a more general level, our analyses emphasize that targets of natural immunity are not polarised between the completely invariant and the highly polymorphic but also include a range of epitopes of intermediate variability that have, so far, been largely overlooked. Among pathogens where protective immune responses are directed against conserved targets or epitopes, a single infection ensures lifelong immunity and the goal of vaccination is to elicit natural immunity against such conserved targets. Most infectious diseases (such as measles) against which we have effective vaccines fall into this category. By contrast, the primary targets of protective immunity in several important pathogens such as *P. falciparum,* are highly diverse and present a challenge to the development of vaccines.

One solution to this problem that is currently being explored by several research groups is the artificial boosting of immunity towards conserved regions that are not sufficiently naturally immunogenic to prevent further infection upon single exposure but do eventually confer protection after multiple exposures. Another strategy is to develop multi-allelic vaccines in which immunogenic epitopes that confer protection against a sub-population of strains are combined to provide broad coverage. However, many of the known targets of protective immunity are too diverse for this strategy to be successfully implemented. Our work in this paper highlights that between these two extremes, there exist epitopes of intermediate variability which are nonetheless potentially under strong immune selection and thus offer alternative targets of vaccination.

## Methods

### Genetic sequence selection

*Pf*AMA-1 sequences were obtained by searching the NCBI Entrez nucleotide and protein databases. This yielded 2835 protein sequences and 2926 nucleotide sequences (up to and including year 2014). In cases when the GenBank nucleotide record did not include the corresponding protein sequence, these were translated using Biopython, an open source set of tools (biopython.org).

The analysis was restricted to sequences between 400 and 622 amino acids in length. The lower limit aims to ensure that all the polymorphic regions across the three domains are included; the upper limit corresponds to the entire processed protein, and therefore removes any entries which contain the non-coding sequence. To avoid pseudoduplication, all identical sequences were removed, leaving 1198 unique sequences. These were then aligned using Clustal W [40].

### Measuring non-overlap among dimorphic sites

A frequency analysis of all residues within the three domains was used to identify any polymorphic sites. Only variants with a frequency exceeding 2.5% were included in the subsequent analysis. This identified 44 dimorphic sites (Figure S1). We searched for signatures of immune selection by analysing the mutual information (MI) scores of all pairs of sites (Table S1).

### Geographic sub-analysis

We repeated the calculation of MI for all isolates whose country of origin could be identified (either from the Entrez database or with reference to published papers).

### Phylogenetic analysis

We first truncated our protein alignment to the region where we had >99% coverage [namely, from residue 149 to residue 534]. This alignment was then used to construct a Maximum Likelihood Tree in Mega6 under the JTT model (with also: 4 discrete categories of gamma-distributed rates among sites; the nearest-neighbour-interchange heuristic; and a very strong branch swap filter) and this tree was used to calculate the maximum parsimony score (PS) for each dimorphic site (Figure S2; Table S1). We defined a new statistic S as the product of the MI and the geometric mean of the PS for each site (Table S2). We used Seq-Gen to simulate 1000 sets of protein sequences with the same phylogenetic characteristics, restricting variation at each of the sites to two amino acids. We were able to establish that the pairs of interest (see Table 1) lay significantly outside the distribution of the identically ranked (by S-score) sites from each simulation (Figure S3).

### Tertiary structures

From the protein data bank (PDB) the sequences and tertiary structures of *Plasmodium vivax* AMA-1 (*Pv*AMA-1, PDB 1W81) and *Pf*AMA-1 domain III (PDB 1HN6) were downloaded. As the tertiary structure of 1HN6 was mostly unstructured, it could not be used on its own. Therefore a Fold and Function Assignment System (FFAS) was used to align the amino acid sequences of 1W81 and 1HN6. From this alignment a ‘hybrid’ structural model with 94% confidence was generated using SCWRL. This resulting model gave a single tertiary structure with the residues within domain III from *Pv*AMA-1 (1W81) substituted by those from *Pf*AMA-1 (1HN6) using the *P. vivax* structural coordinates. Pymol (http://www.pymol.org/) was used to visualise the ‘hybrid’ tertiary structure and to highlight the relevant residues.

## Supplementary files

Table S1: parsimony score and MI for pairs of sites

Table S2: statistic S (product of MI and the geometric mean of the PS for each site)

## Acknowledgements

We thank Dr. CRC Spensley for assistance with data manipulation and programming, and Dr. Pietro Roversi for assistance with generating the structure.

## Author Contributions

JL, KJS, PSW, SG designed the study. JL, PSW, AW performed the experiments. All authors wrote and revised the manuscript. JL, KJS, PSW curated the data.

